# Decoding Parametric Grip-Force Anticipation from fMRI-Data

**DOI:** 10.1101/2024.09.19.613884

**Authors:** Guido Caccialupi, Timo Torsten Schmidt, Till Nierhaus, Sara Wesolek, Marlon Esmeyer, Felix Blankenburg

## Abstract

Previous functional magnetic resonance imaging (fMRI) studies have shown that activity in premotor and parietal brain-regions covaries with the intensity of upcoming grip-force. However, it remains unclear how information about the intended grip-force intensity is initially represented and subsequently transformed into a motor code before motor-execution. In this fMRI study, we used multivoxel pattern analysis (MVPA) to decode where and when information about grip-force intensities is parametrically coded in the brain. Human participants performed a delayed grip-force task in which one of four cued levels of grip-force intensity had to be maintained in working memory (WM) during a 9-second delay-period preceding motor execution. Using time-resolved MVPA, with a searchlight approach and support vector regression (SVR), we tested which brain regions exhibit multivariate WM codes of anticipated grip-force intensities. During an early delay-period, we observed above-chance decoding in the ventromedial prefrontal cortex (vmPFC). During a late delay-period, we found a network of action-specific brain regions, including the bilateral intraparietal sulcus (IPS), left dorsal premotor cortex (l-PMd) and supplementary motor areas (SMA). Additionally, cross-regression decoding was employed to test for temporal generalization of activation patterns between early and late delay-periods with those during cue presentation and motor execution. Cross-regression decoding indicated temporal generalization to the cue-period in the vmPFC, and to motor-execution in the l-IPS and l-PMd. Together, these findings suggest that the WM representation of grip-force intensities undergoes a transformation where the vmPFC encodes information about the intended grip-force, which is subsequently converted into a motor code in the l-IPS and l-PMd before execution.

## 1 INTRODUCTION

Imagine a mountain bike rider steering down a challenging section of a gravel hill slope. As they recognize an impending transition to steeper and firmer terrain, they decide to reduce the bicycle’s speed by applying a grip-force to the handlebars. However, to avoid a severe fall on the unstable terrain, the bike rider must prepare and delay the grip-force application. This scenario illustrates the continuity of serial translational processes, from the selection of an intended action to the motor planning of a grip force application through working memory (WM) maintenance, to motor execution. Until today, there has been substantial research on the question of how a specific action is selected (Soon *et al*., 2008, 2013; Bode & Haynes, 2009). However, it remains challenging to thoroughly characterise the translational processes from action selection to motor execution, in particular when an abstract goal is transformed into a motor movement plan (Kim *et al*., 2021). Therefore, systematically testing where in the human brain an intended action is represented during an early phase (after action selection) as compared to a later period (closer to motor execution) can provide further insight into this process.

The application of decoding techniques to functional magnetic resonance imaging (fMRI) data has provided a powerful tool for investigating where information is represented in the human brain (Haynes & Rees, 2006; Norman *et al*., 2006). In particular, the use of multivoxel pattern analysis (MVPA) in delayed response paradigms (*e.g.*, Harrison & Tong, 2009) has proven successful for testing where and when information is encoded during different processing phases of WM (Schmidt *et al*., 2017; Hebart *et al*., 2018). Of particular interest in WM studies are the transformation phases from the initial presentation of a stimulus (or task cue) to its representation during the early and late phases of a delay period. Previous research has shown that stimulus information is represented in sensory areas during and immediately after the stimulus presentation in visual (Christophel *et al*., 2016), tactile (Schmidt *et al*., 2021), and auditory (Uluç *et al*., 2018) WM studies. These studies further describe how information is transformed into more abstract representations in higher-order sensory and multimodal cortices depending on task demands (Christophel *et al*., 2017). While these studies focus on the WM delay period, they do not examine how this information is translated into a motor plan for execution (Bode & Haynes, 2009). In more ecologically valid scenarios, however, the primary function of WM is assumed to be using information to act on the environment (Parr & Friston, 2017). As a result, recent research has emphasized the importance of studying how WM information is transformed into specific action plans (*e.g.*, Nobre & van Ede, 2023). To date, only a few studies have investigated how information is encoded when a specific motor plan is cued, maintained, and then executed after a delay period (*e.g.*, Boettcher *et al*., 2021).

Recent fMRI MVPA studies have shown that action-specific information, such as the outcome of an intended action (or its “goal”; Hamilton & Grafton, 2007, 2008; Turella *et al*., 2020), can be decoded from the ventromedial prefrontal cortex (vmPFC) during an early delay period, even before motor planning begins (Soon *et al*., 2008, 2013; Ruiz *et al*., 2024). This finding aligns with a two-stage model in which information about the intended action is first selected, and then, motor movements are specified and prepared for execution (Boettcher *et al*., 2021). This notion aligns with findings from transcranial magnetic stimulation (TMS) and fMRI decoding studies (Tecilla *et al*., 2022), which suggest a distinction between a motor decision phase in which a motor plan is selected from alternative actions, and a subsequent movement planning phase (Nakayama *et al*., 2008; Tecilla *et al*., 2022; Ruiz *et al*., 2024), with distinct brain regions being recruited during each phase. The above-chance decoding in the vmPFC is also consistent with findings from a recent EEG study (Boettcher *et al*., 2021), which showed that in the absence of sensory stimulation, the electrophysiological brain activity reflects the outcome of an intended action before the instruction of movements (Ruiz *et al*., 2024) or selection of motor parameters (*e.g.*, specific muscle fibres; Mizuguchi *et al*., 2011, 2013, 2014). Collectively, these studies highlight a distinction between the initial motor-decision stage and the subsequent planning stage, emphasizing that motor planning does not commence until the intended action is selected.

After action selection, motor planning occurs (Nakayama *et al*., 2008; Tecilla *et al*., 2022; Ruiz *et al*., 2023) and is conceived as a neural process that optimizes an ideal preparatory state for motor output generation (Vyas *et al*., 2020; Churchland *et al*., 2010; Shenoy *et al*., 2013; Ariani *et al*., 2022). Animal studies on motor planning, which employed single-cell recordings, have consistently found preparatory signals in patterns of neuronal firing in the dorsal premotor cortex (PMd; *e.g.*, Cisek & Kalaska, 2004, 2010; Hoshi & Tanji, 2006, 2007), the supplementary motor area (SMA; *e.g.*, Hoshi & Tanji, 2004), and the PPC (*e.g.*, Cui & Andersen, 2007; Cui & Andersen, 2011; Andersen & Cui, 2009). Human fMRI MVPA studies have shown that activity in parieto-frontal brain regions during the planning of grasping actions was predictive of several movement properties, such as the kinematics of grip types (Gallivan *et al*., 2011b; Ariani *et al*., 2015; Ruiz *et al*., 2024), the action order (Gallivan *et al*., 2016; Ariani *et al*., 2022), and the to-be utilized effector (Gallivan *et al*., 2011a, 2013; Leoné *et al*., 2014; Turella *et al*., 2016). Delayed grip-force tasks have also been used to investigate the neuronal underpinnings of the anticipatory scaling of grip-force intensities (Cole & Rotella, 2002; Chouinard *et al*., 2005; Nowak *et al*., 2009; van Nuenen *et al*., 2012). In particular, fMRI studies using univariate analyses have shown that the strength of the blood-oxygen-level-dependent (BOLD) signal in the PMd and PPC reflects the anticipated grip-force intensity (van Nuenen *et al*., 2012; Mizuguchi *et al*., 2014).

To isolate neural correlates of motor planning from execution, previous studies have used delayed-movement tasks, which resemble the delayed-match-to-sample (DMTS) or categorization (DMTC) tasks commonly used in working memory research. These include delayed grip-force tasks (*e.g.*, van Nuenen *et al*., 2012) that, requiring the maintenance of a graded motor parameter over time, are highly similar to parametric WM tasks (*e.g.*, Romo *et al*., 1999; Brody *et al*., 2003). Romo and co-workers revealed that maintained continuous sensory parameters, such as vibrotactile stimulation intensities, are encoded in persistent, monotonically tuned neural activity in the (right) inferior prefrontal cortex (*e.g.*, Romo *et al*., 1999; Brody *et al*., 2003). More recently, MVPA has been essential in showing that premotor and parietal brain regions can parametrically represent and maintain information in distributed patterns of brain activity (Schmidt *et al*., 2017; Uluç *et al*., 2018; Wu *et al*., 2018; Uluç *et al*., 2020). However, it remains to be tested whether parametric information about grip-force intensities is also encoded in multivariate patterns during action selection and motor planning (*i.e.*, during the early- and late delay periods).

The present study investigates whether, where and when grip-force intensities can be parametrically decoded from fMRI data in a delayed grip-force task. To relate our findings to the parametric WM findings, we modified the established DMTS to a delayed grip-force task. However, instead of maintaining sensory properties of stimuli such as vibrotactile frequencies (*e.g.*, Romo *et al*., 1999; Schmidt *et al*., 2017), participants had to anticipate and maintain grip-force intensities before execution. Thereby, we decoded parametric grip-force intensities from the early- and late delay periods using time-resolved MVPA (by employing support vector regression; SVR). In addition, cross-regression decoding was used to test whether neural representations during the early- and late delay-periods were similar to those during the motor execution and cue periods (Myers *et al*., 2008; King & Dehaene, 2014; Hebart & Baker, 2017; Hebart *et al*., 2018; Ariani *et al*., 2015; Ariani *et al*., 2018; Nierhaus *et al*., 2023). Based on the literature, we hypothesized that parametric neural representations of grip-force intensities can be identified in action-specific brain regions, such as the PMd, SMA and PPC, and that these codes differ from early representations in the vmPFC.

## 2 MATERIALS AND METHODS

### 2.1 Participants

All participants (*N* = 33) were healthy and right-handed, as assessed by the *Edinburgh Handedness Inventory* (*EHI;* Oldfield, 1971). Before the experiment, they provided written informed consent to participate in the study in accordance with protocols approved by the local ethics committee of *Freie Universität Berlin* (003/2021, Berlin, Germany).

To account for potential biases and confounds, only *N* = 22 participants were included in the analysis (age: 30 ± 6.31, 4 female). Five participants were excluded due to low performance (*i.e.,* percentage of trials with accurate responses < 25% in at least one experimental run and one experimental condition). Another six participants were excluded due to frequent grip-force application during the delay period (*i.e.*, percentage of trials with applied force > 75% in at least one run and one condition). To further ensure that the analyses were not confounded by early motor responses, we excluded from the dataset all trials in which any grip-force was applied during the delay period (cut-off: mean grip-force ≥ 0.05 on a scale from 0 to 1).

### 2.2 Experimental procedure

After participants entered the MRI scanner, the grip-force transducer was calibrated to participants’ maximum grip-force. Next, participants were introduced to the four grip-force levels that served as the target grip-force intensities in the main experiment. In this training phase, they were familiarized with the grip-force transducer by performing twenty-four trials with real-time visual feedback of the applied grip-force. In twenty-four further trials, they were trained on the experimental task (see below) during the acquisition of a structural MRI scan. Finally, participants performed the delayed grip-force task (illustrated in **Figure 1A**) in four runs during fMRI scanning.

**Figure 1.**
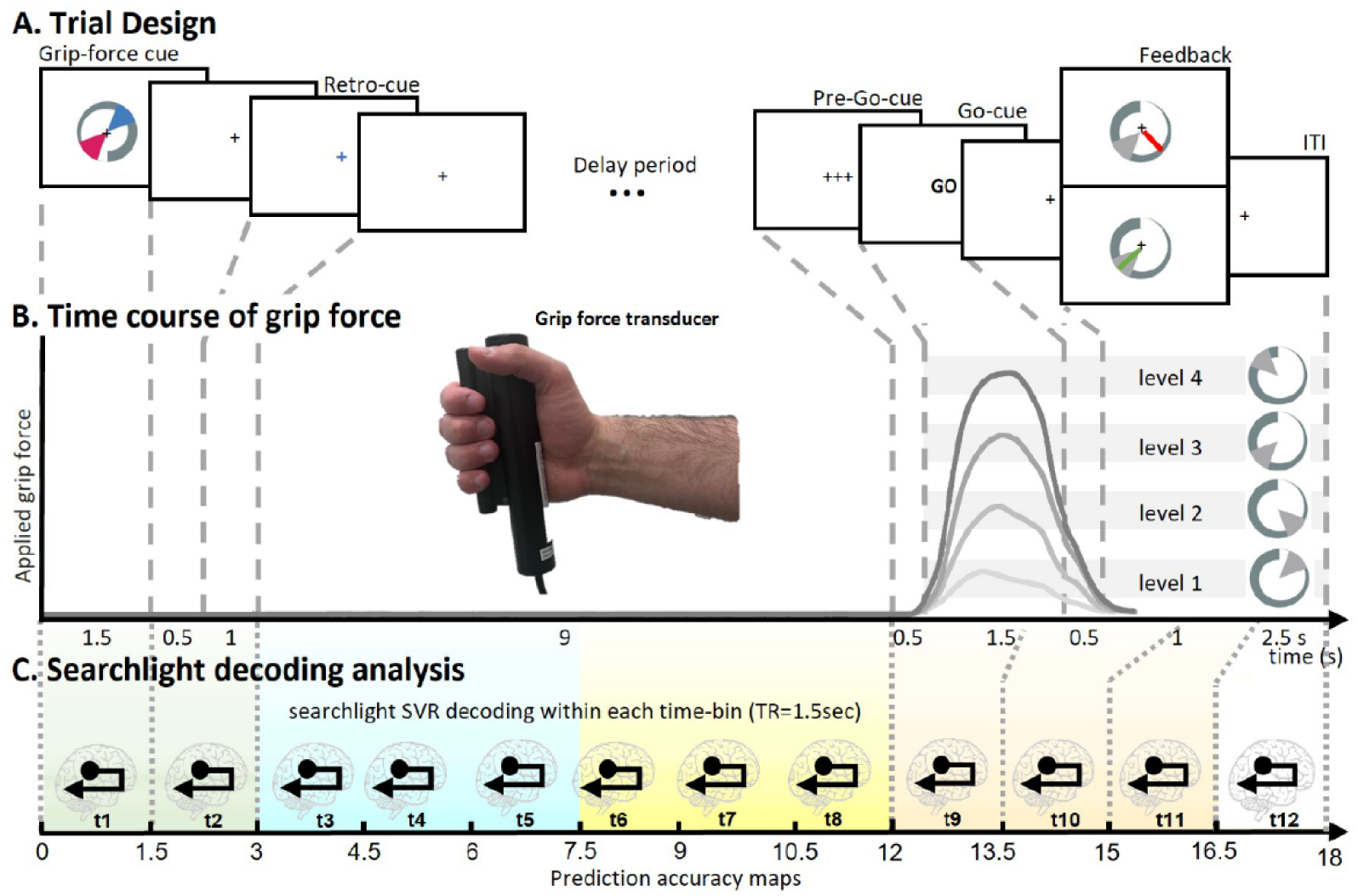
**A**. *Delayed grip-force paradigm*. Two grip-force levels were presented on a *grip-force cue (i.e.*, a grey circular increasing bar with one level represented in cyan and the other in red, as sectors of 50° on the *indicator*). A *visual retro-cue* (*i.e.*, a cyan or red cross displaced in the centre of the screen) indicated which grip-force had to be anticipated and maintained during a subsequent 9 s *delay period*. After this *delay*, participants prepared and performed the grip force upon the presentation of a *pre-Go* (0.5 s) and a *Go-cue* (1.5 s). Subjects responded with their right hand by differently squeezing a grip-force device, and a *Feedback* representing the applied force was provided at 1.5 s from the *Go-cue*. **B**. Different shades of grey represent time courses of mean grip forces applied during the experimental-trial, calculated over the accurate trials per condition *(i.e.*, per grip-force level). Light-grey bars displayed in the background represents grip-force levels. **C**. Represents the multivoxel pattern analysis (MVPA) with a searchlight and a time-resolved approach adopted across four time periods of the trial. The pastel green background represents a 3 s cue period, the light cyan background depicts a 4.5 s early delay period, the light yellow background refers to a 4.5 s late delay period, and the light red background marks a 4.5 s motor execution period.

### 2.3 Grip-force assessment and stimuli

Based on the visual presentation of a *grip-force cue* and a *retro-cue*, participants had to apply a grip force via the right hand on a cylindrical MR-compatible grip-force transducer (*i.e.*, a *force fibre optic response pad*, Current Designs, HHSC-1x1-GRFC-V2; illustrated in **Figure 1B**). Grip-force intensity was sampled throughout the task to ensure that participants only applied force during the execution period of the trials, allowing the exclusion of trials where participants applied force during other periods of the experimental trials.

The visual *grip-force cue* was presented in the centre of the screen, utilizing *Psychtoolbox-3* (Brainard, 1997). The visual cues were composed of a 360° circular display increasing in thickness, corresponding to an increase in grip-force where the maximum corresponded to 75% of the individual maximum grip-force. Four grip-force levels were indicated by the display of a range, defining a sector of 50° of the grip-force spectrum, with grip-force level 1: 20°-70° (5-20% of maximum grip force level), level 2: 110°-160° (30-45%), level 3: 200°-250° (55-70%), and level 4: 290°-340°(80-95%) (as illustrated in **Figure 1B**, right side; and **Figure 2B**). The *grip-force cue* was presented for each trial with a random degree of rotation (as illustrated in **Figure 1A**).

**Figure 2.**
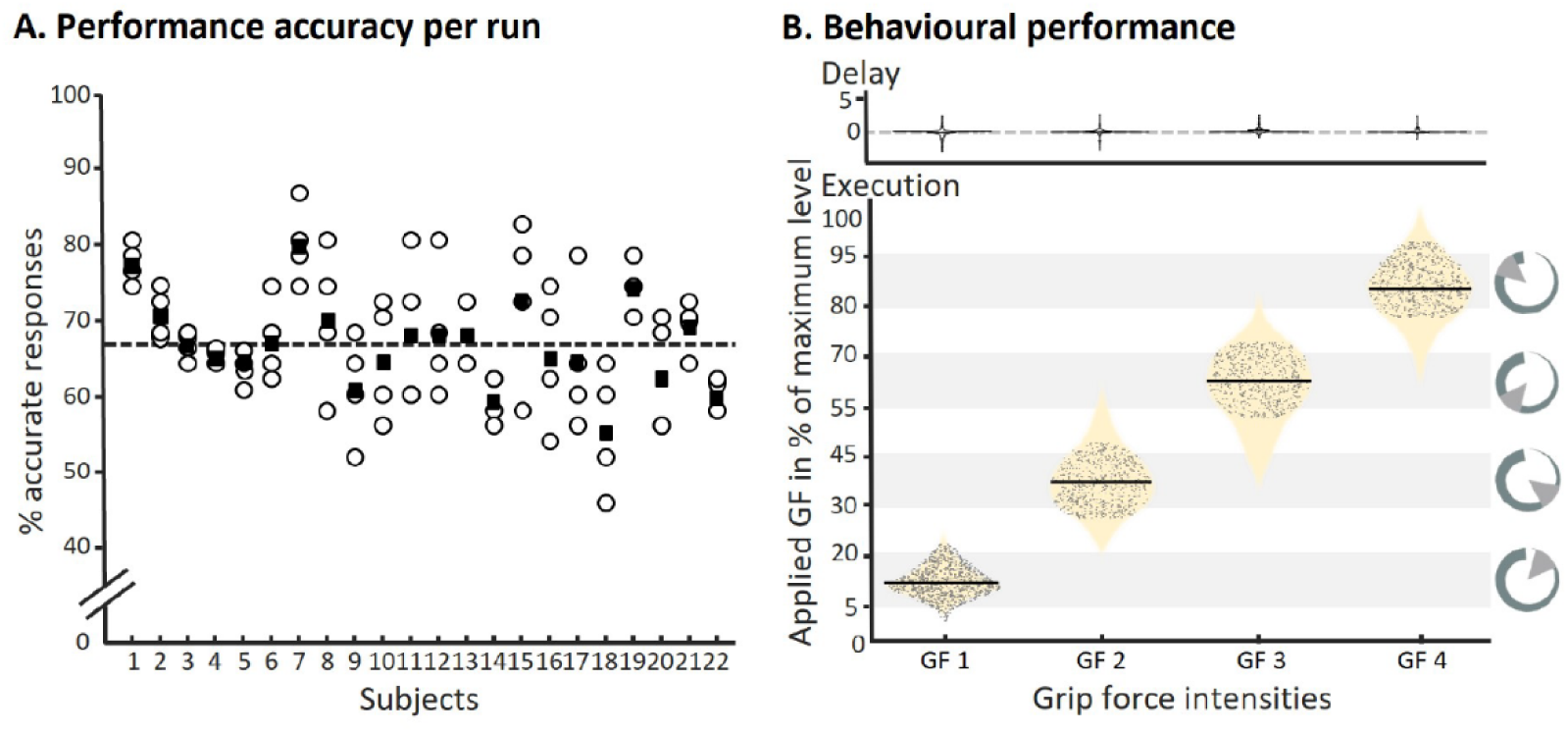
Behavioural assessment. **A.** Shows that participants performed consistently across all four runs of the delayed grip- force task. Circles represent the performance of accurate grip-force application in each of the four runs (for each participant); filled squares represent the mean performance across runs. The overall mean performance across participants is represented by the dashed black line. **B.** Displays the grip-forces applied during the delay period (upper display) and the execution period (lower displays). We made sure that only trials were included in the analysis, where participants did not press the grip-force device during the delay period (upper panel), and accurate grip-force was considered within the target level ± 20° (lower panel). Light-grey bars in the background represent the cued grip-force (GF) levels, where fore each of the four cued GF levels a violin plot (beige) represents the distributions of applied grip-forces, with the trials included in the analysis indicated as black dots.

### 2.4 Experimental task

During fMRI scanning participants performed a delayed grip-force task. Each trial started with the presentation of a *grip-force cue*, comprised of two grip-force levels presented in cyan and red (illustrated in **Figure 1A**). A *retro-cue* indicated which of the two grip-force intensities had to be maintained and executed. The combinations of displayed grip-force levels were balanced so that all combinations of two different levels were presented equally often. After a *delay period* of 9 seconds, participants had to apply the anticipated grip-force upon the display of a *Go-cue*; importantly, in the absence of the grip-force cue. A *pre-Go-cue* was presented for 0.5 s before the response to allow optimal response timing during the 1.5 s response window. Finally, visual *Feedback* was provided by showing on a screen the applied force as a green radian if accurate or as a red radian if inaccurate, *i.e.* outside of the cued grip-force level (compare **Figure 1A**). The mean grip-force during the last 0.75 s of the response window was evaluated to assess accuracy of responses. For the analysis, trials were considered accurate when the applied force was within the target range ± 20° (5% of maximum grip force).

Four experimental runs comprised 48 experimental trials supplemented with 12 catch trials with a shorter delay period of 6.0 s, 4.5 s, 3.0 s or 1.5 s. Catch-trials were designed to enhance participants’ readiness to prepare and execute the grip-force task, and thus the continuous maintenance of the force intensity. Catch trials were not included in the analyses. Trial order was fully randomized within a run, where each experimental condition (*i.e.*, the cueing of each of the four to-be anticipated grip-force intensities) was presented equally often (*i.e.*, 12 times per run).

### 2.5 fMRI data acquisition and preprocessing

fMRI data were acquired in 4 runs of 18 min and 40 s each on a 3T Siemens Prisma at the *Center for Cognitive Neuroscience Berlin* (CCNB) of the *Freie Universität Berlin*. For each run, 745 functional images were obtained with an EPI sequence (64 channel head coil, 48 slices, interleaved order; TR= 1.5 sec, 2.5 × 2.5 × 2.5 mm voxel size; multiband acquisition with acceleration factor of 3).

Trial onsets were time-locked to the functional image acquisition to allow a time-resolved analysis (see below). MRI data processing was performed using SPM12, r7771 (*Wellcome Trust Centre for Neuroimaging*, *Institute for Neurology*, *University College London*). To preserve the spatiotemporal structure of the fMRI, data preprocessing was limited to spatial realignment.

#### 2.5.1 Time-resolved searchlight decoding of grip-force anticipation

We applied an MVPA searchlight approach to identify brain regions that exhibited multivariate parametric codes of grip-force anticipation during the delay period. Finite impulse response (FIR) models were used to obtain run-wise beta estimates for each 1.5 s time-bin of the trials. 12 consecutive time-bins were modelled (as illustrated in **Figure 1C**), comprising 6 time-bins throughout the delay period, 2 additional time-bins before the delay period and 4 time-bins from grip-force execution. High-pass filtered data (cut-off 128 s) were included in a corresponding first-level GLM model with 220 regressors (4 conditions × 12 time-bins × 4 runs complemented with the 6 realignment parameters). To balance the number of trials included in each condition, allowing equally reliable beta-estimates, we subsampled the trial number to N = 5 for condition (including the trials with most precise grip-force performance).

To identify where and when information about the maintained grip-force was encoded in the brain, we applied time-resolved MVPA using SVR (Kahnt *et al*., 2011), which can be considered a multivariate pendant of a parametric coding, a method previously applied in a series of WM decoding experiments (*e.g.*, Christophel *et al*., 2012; Schmidt *et al*., 2017; Uluç *et al*., 2020 Pennock *et al*., 2021). Applying a searchlight approach (r=4 voxel) independently for every time-bin allows testing for local multivariate representations (*i.e.*, activation patterns of voxels) that code parametric grip-force anticipations. An SVR model is trained to predict grip-force (*i.e.*, the four grip-force intensities) based on a multivariate data vector (*i.e.*, multivoxel activation pattern). This is similar to a univariate regression approach, where prediction of the dependent variable is however solely based on one independent variable (see Kahnt *et al*., 2011 for more information).

To account for differences in the difficulty of grip-force performance, the distances between the four SVR labels were adjusted using Fechner’s law (Fechner, 1860). First, we accounted for differently perceived difficulties by log-transforming the applied grip-force intensities, which were grouped in four normal distributions of applied grip-force. After, we calculated the three distances between the four labels of the SVR (*d_i_*, *d_i_*_+1_, *d_i_*_+2_). Specifically, such distances were obtained by calculating the inverse of the difference between the means for each combination of increasing grip-force intensities 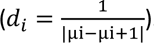. We used the resulting distances to adjust the SVR’s labels (with resulting labels: −1.5, −0.58, 1.36, and 4.33).

Beta estimates of each condition were first normalized (*i.e.*, z-scaled) across the samples for each voxel as implemented in TDT (Hebart *et al*., 2015) and forwarded to a four-fold leave-one-run-out cross validation schema. To make our data comparable to previous reports, we used as measure the prediction accuracy defined as the Fisher’s z-transformed correlation coefficient between the predicted value levels and the actual value levels of the test dataset (in TDT: “zcorr”, see also Kahnt *et al*., 2011). The centre of the searchlight was moved voxel wise through the brain, and prediction accuracy values were saved as corresponding whole-brain accuracy maps. In this way, we obtained an accuracy map for every subject and time-bin, reflecting local activation patterns that code grip-force intensity in a multivariate way.

Prediction accuracy maps were entered into a second-level analysis to test for above-chance decoding in terms of SPM’s flexible factorial design implementation of an ANOVA. We used t-contrast to test for above chance decoding across different time-bins. All results are reported at a threshold of p < 0.05 family-wise error (FWE) corrected.

#### 2.5.2 Control analysis: decoding the non-memorized grip-force

To test for the specificity of the main analysis, we performed a second MVPA as a control analysis. Namely, we tested for above-chance prediction accuracy for the non-maintained grip-force intensities. A new FIR model was estimated, in which four sets of FIR regressors modelled those trials in which a grip-force level was presented and not retro-cued. As previously done in the main analysis, 12 consecutive time-bins were modelled for the 5 most accurate trials of each condition. High-pass filtered data (cut-off 128 s) were included in a corresponding first-level GLM model with 220 regressors (4 conditions × 12 time-bins × 4 runs complemented with the 6 realignment parameters). Therefore, each beta image was estimated with equal amounts of data (*i.e.*, modelling the same number of trials) as in the main analysis. Beta images were entered into an identical SVR searchlight and second-level analysis as in the main analysis.

#### 2.5.3 Test for temporal generalization: Cross-regression decoding

To test whether the multi-voxel activation patterns in the early and late delay periods were similar to those in the cue and motor execution periods, we conducted a cross-regression decoding analysis (as test for temporal generalization). We use the term “cross-regression” instead of “cross-classification” because a linear SVR model was trained to predict a continuous variable (*i.e.*, grip-force intensity) rather than to classify into a category (Kahnt *et al*., 2011; Schmidt *et al*., 2017). Nonetheless, the logic underlying the use of cross-classification to test for temporal generalization applies to SVR approaches as well (Bode *et al*., 2022). Similar to previous MVPA studies (Myers *et al*., 2008; King & Dehaene, 2014; Hebart *et al*., 2018), we tested for temporal generalization by training on each time-bin and respectively testing on all other time-bins. The cross-regression analysis was based on the same data as the main analysis, namely: the beta estimates derived from the same FIR model were used. SVRs was trained and tested on all time bins (t1-t12 x t1-t12) using four-fold leave-one-run-out cross-validation as in the main analysis, resulting in 144 cross-regression accuracy maps.

For the statistical assessment, we averaged prediction accuracy maps to fall into 16 cross-regression accuracy maps to reflect the 4 time-periods investigated in the main decoding analysis, *i.e.*, the cue period, early delay period, late delay period, and motor execution period. After normalization and smoothing, the 16 cross-regression prediction accuracy maps of 22 participants were entered into a second-level analysis to test for above-chance decoding using SPM’s flexible factorial design specification of an ANOVA. This design included one factor with four levels for the training periods, and a second factor with four levels for the testing periods. We used t-contrasts to assess above-chance cross-regression decoding on each of the 16 maps. All results are reported at *p* < 0.05 FWE corrected.

## 3 RESULTS

### 3.1 Behavioural performance

The participants included in the main analysis (N = 22) responded with accurate application of grip-force (*i.e.*, they were in the target range ± 20°) in 67 ± 5.5% (mean ± SD) of the trials. Average performance across runs indicates small training effects over time: run 1: 66 ± 7.7%; run 2: 64 ± 7%; run 3: 70 ± 10%; run 4: 70 ± 7% (performance per participant and run are illustrated in **Figure 2A**). When tested with a 1 x 4 repeated-measures ANOVA, this effect of runs was significant (*df* = 3, F= 2.73, *p* = 0.0491, η^2^ = 0. 0178), while post-hoc t-tests did not reach significance after Bonferroni correction. These findings render major learning effects on the neuroimaging results rather unlikely.

To test for performance differences across the four to-be-applied grip-force intensities, we performed a 1x4 repeated-measures ANOVA, which was significant (*df* = 3, F= 78.35, *p* = 0.001, η^2^ = 1. 8017). This was also reflected by significant post-hoc t-tests, which revealed that the application of grip-force intensity 1 was easier than the other grip-force intensities. After Bonferroni correction, the mean of the performance accuracy for grip-force intensity 1 was significantly greater than intensity 2 (M_diff_ = 26%, *p* = 0.001), intensity 3 (M_diff_ = 31%, *p* = 0.001), and intensity 4 (M_diff_ = 26%, *p* = 0.001). Violin plots in **Figure 2B** display the distribution of responses (*i.e.*, accurate and non-accurate performances) in terms of applied force on the grip-force device for the respective trial types (*i.e.*, grip-force intensities); the data points display only accurate responses. We further ensured that the included participants did not apply force during the *delay period* (see **Figure 2B**, upper display). These results are in line with an expected and previously reported Fechner-Law effect for perceived difficulties of to-be executed force intensities (Jones, 1989). Therefore, we applied a correction to the fMRI data modelling (see *Methods* section).

### 3.2 Multivariate mapping of regions that code grip-force anticipation

To identify brain regions that encode parametric grip-force anticipation, we conducted a time-resolved whole-brain SVR analysis. Within a second-level ANOVA design, we computed t-contrasts across prediction accuracy maps of 3 periods of interest, *i.e.*, the cue period (C, time-bins t1-t2), the early delay period (ED, t3-t5) and the late delay period (LD, t6-t8); all *p* < 0.05 FWE-corrected. No brain region exhibited above-chance decoding during the cue period, only a small cluster in the vmPFC was revealed when assessed at *p* < 0.001 uncorrected. During the early delay period, a cluster in the ventromedial prefrontal cortex (vmPFC) was revealed. In the late delay period, a network including bilateral intraparietal sulcus (IPS), left dorsal premotor cortex (l-PMd), and the supplementary motor area (SMA) was found (see **Figure 3A**, **Table 1**). Finally, a t-contrast on the motor execution period (*i.e.*, ME, t9-t11) revealed one extended cluster spanning bilateral primary motor cortex (M1) and primary somatosensory cortex (S1), parietal cortices, and the cerebellum (**Figure 3A**, fourth column, and **Table 1**).

**Figure 3.**
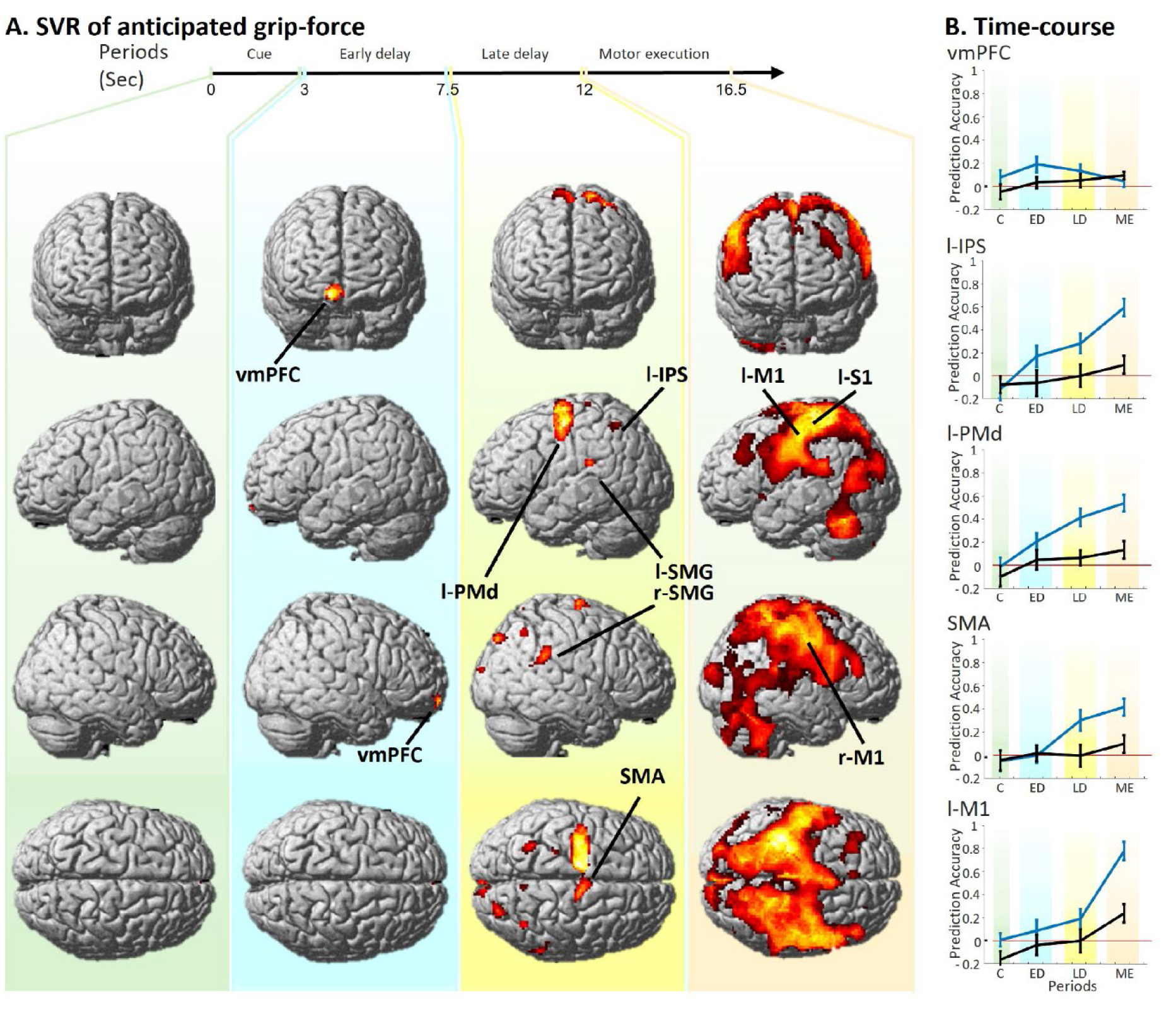
Results of time resolved support vector regression analysis **A.** Displays brain regions that parametrically code grip-force intensities during the cue period (C), early delay period (ED), late delay period (LD) and motor execution period (ME). Brain regions with above-chance prediction accuracy are revealed by four t-contrasts displayed at *p* < 0.05, FWE-corrected. No brain region exhibited above-chance decoding during C (as displayed in the first column, pastel green background). During the early delay period, a cluster in the vmPFC was revealed (second column, light cyan background). In the late delay period, a network including bilateral intraparietal sulcus (IPS), left dorsal premotor cortex (l-PMd), and the supplementary motor area (SMA) was found (third column, light yellow background). Finally, a t-contrast on the motor execution period revealed one extended cluster centred on the M1 and S1 (fourth column; light red background) **B.** Time-courses of prediction accuracy values relative to the main decoding analysis (displayed in blue) and the control analysis (in black). Prediction accuracy values were extracted from the peak voxels of the five most representative clusters reported in Table 1. The time-course represents four prediction accuracy values obtained by averaging prediction accuracy values of twelve time-bins in correspondence of the four time-periods (tested in the main analysis). As expected, prediction accuracy values for the control analysis do not overcame chance level. The control analysis thereby demonstrates the specificity of the main decoding analysis for the maintenance of grip-force intensities.

**Table 1.** Regions that exhibit above-chance prediction accuracy across the cue period, early and late delay periods and motor execution period, revealed by a t-contrast displayed at *p* < 0.05, FWE corrected.

| Cluster size | Anatomical Region | Peak MNI coordinates |  |  | z-score |
| --- | --- | --- | --- | --- | --- |
|  |  | x | y | z |  |
| Early delay Period |  |  |  |  |  |
| 231 | Ventromedial PFC | 6 | 60 | -10 | 5.27 |
| Late delay Period |  |  |  |  |  |
| 1370 | Left PMd | -26 | -10 | 58 | 6.48 |
|  | SMA | -6 | -6 | 46 | 5.16 |
| 156 | Right SMG | 52 | -42 | 32 | 4.94 |
| 73 | Left IPS | -30 | -54 | 52 | 4.89 |
| 42 | Right IPS | 38 | -58 | 48 | 4.74 |
| Motor execution Period |  |  |  |  |  |
| 58068 | Left S1 | -32 | -34 | 58 | 11,42 |
|  | Left M1 | -43 | -14 | 48 | 8.46 |

To plot the temporal evolution of grip-force specific decoding accuracy within the network of brain regions recruited during the early and late delay periods, we extracted the time-course of prediction accuracies from the peak voxels of the identified clusters in the vmPFC, l-IPS, l-PMd, SMA and l-M1 (see **Table 1** for MNI coordinates). Time-courses of mean prediction accuracies within the four time-periods of the experimental trials are displayed in **Figure 3A**. As expected from the dynamics of the BOLD response and previous reports (*e.g.*, Schmidt *et al*., 2017; Uluç *et al*., 2020), the prediction-accuracies peaked between 4 to 8 s after the start of the delay period (see also *Supplementary Materials* for time-courses plotted for 12 time-bins). The time-course of the vmPFC displayed an early peak during the early delay period, preceding those of action-specific brain regions, which reached maximum in the late delay period. All regions of interest, except the vmPFC, showed a substantial increase of prediction accuracy during the execution.

### 3.3 Control analysis: decoding non maintained grip-force intensities

As a control analysis for the specificity of the main analysis, we tested if information on the non-memorized (*i.e.*, non-cued) grip-force intensity can be decoded. Therefore, we conducted an identical MVPA as the main analysis (based on FIR models that modelled the non-cued intensity). Please note that the amount of cued and non-cued grip-force intensities was balanced within and across runs. The decoding analysis of the non-maintained grip-force intensity did not reveal any significant cluster (*p* < 0.05 FWE-corrected), corroborating the specificity of the main analysis. Time-course data of this control analysis is displayed in **Figure 3B** and in the *supplementary materials*.

The specificity of the main analysis is further corroborated by the results of a label permutation test (*i.e.*, a supplementary control analysis), which indicated that prediction accuracies of the main analysis are indeed based on the parametric coding of grip-force intensity anticipation. Additionally, results of two univariate control analyses rendered unlikely that parametric effects found in the main MVPA analyses were mostly driven by univariate effects (see *Supplementary Materials*).

### 3.4 Testing for temporal generalization

To test whether multivariate patterns of grip-force intensities found in the early and late delay periods were similar to those in the cue and motor execution periods, we conducted whole-brain cross-regression decoding analyses. All combinations of forward and backward generalization were tested (see **Figure 4A**). This means that in forward generalization all combinations of training on early time-bins and testing on later time-bins were computed to test if the WM representation is similar to an initially formed mental representation during the action selection. In contrast, backward generalization trains on the activation patterns during motor execution and tests on the preceding time-bins to evaluate if during the WM period (*i.e.*, anticipation of motor execution) already shows neuronal activation that is similar to the actual motor execution.

**Figure 4.**
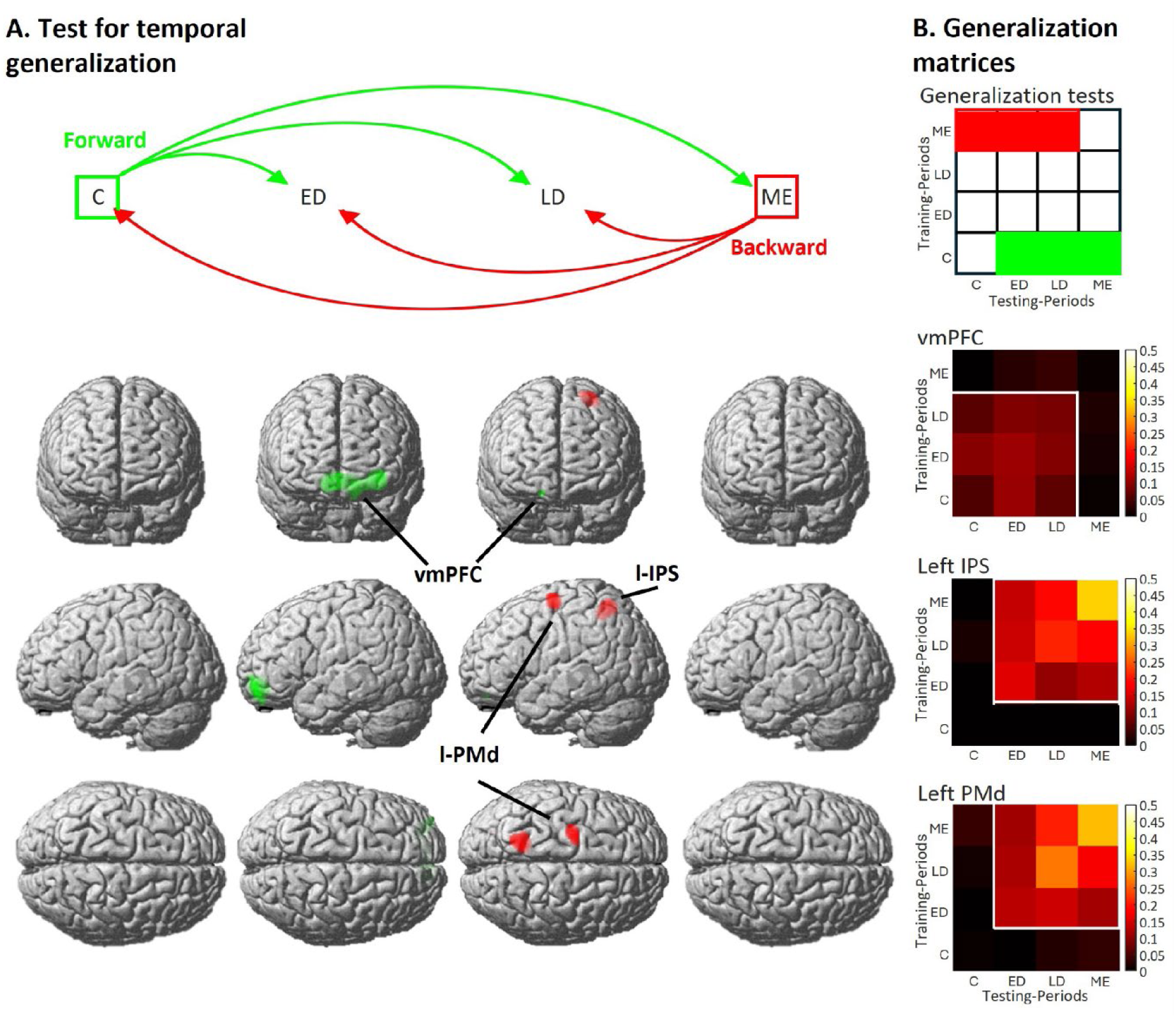
**A.** The upper display shows four time periods: the cue period (C), early delay period (ED), late delay period (LD), and motor execution period (ME). The three green arrows illustrate forward generalization (where the SVR was trained on the C and tested, respectively, on the ED, LD, and ME); red arrows illustrate backward generalization (with training on the ME and testing on the LD, ED, and S). Six t-contrasts were computed on the resulting cross-regression accuracy, and results are showed in the lower display in four columns, corresponding to the testing periods. Results are colored as forward (green) and backward (red) generalization. **B.** The upper display indicates which fields of a generalization matrix correspond to forward and backward generalizations (as displayed in A). The generalization matrix represents testing periods on the *x-axis and* training periods on the *y-axis.* According to this schema, the lower display shows three temporal generalization matrices, each corresponding to a brain region that showed above-chance decoding. These matrices display cross-regression accuracy values for the peak-activity voxel in the vmPFC, l-IPS, and l-PMd (as identified in the main decoding analysis). Sixteen cross-regression accuracy values (reflecting the full range of tests across time-periods; see Method section) are represented within each heatmap. Brighter colours represent higher cross-regression accuracy values. The contours represent above-chance cross-regression decoding, as revealed by t-contrasts on the peak voxel. Overall, the three generalization matrices indicate that prediction accuracy values are similar for both generalization directions (*e.g.*, whether training is conducted on S and testing on ED, or vice versa). All brain regions exhibit relatively stable brain activity patterns between the two delay periods (*i.e.*, when generalization tests are conducted between ED and LD).

First, to test for forward generalization, we computed a t-contrast across the accuracy maps from training on the cue period (C) and testing in the other periods. Results are displayed in **Figure 4A** in green (all results are displayed at *p* < 0.05 FWE-corrected). Interestingly, a single cluster in the vmPFC was revealed, which is largely overlapping with the vmPFC of the main analysis. This cluster was found less pronounced in the late delay period (LD). Consistently, **Figure 4B** shows the stability of generalization when training on the early delay period (ED) and testing on the late delay phase for the vmPFC cluster. In contrast, no temporal generalization was found to motor execution even at *p* < 0.001, uncorrected, indicating that the initial mental representation might be transformed into a different code for motor execution.

Analogously, we tested for backward generalization by training SVRs on the motor execution period (ME) and testing on the preceding periods. Results are displayed in **Figure 4A** in red. Interestingly, only two clusters in the l-PMd and l-IPS were found, which largely overlap with those exhibited in the late delay period of the main decoding analysis (Compare **Figure 3A)**. Also, this analysis did not reveal any temporal generalization to the cue period (even at *p* < 0.001, uncorrected). Thereby, forward and backward temporal generalization tests corroborate that over time the neural representation of the intended action is converted to a different mental representation. Early codes include the vmPFC, while later in the trial information is found in the l-IPS and l-PMd, indicating similar codes as during motor execution.

## 4 DISCUSSION

In this fMRI study, we employed a delayed grip-force task and time-resolved MVPA to identify brain regions where information about grip-force intensities is parametrically coded and transformed during the cue period, early and late delay periods, and motor execution.

As expected, information about the intended grip-force intensity could only be decoded starting from the delay period (*i.e.*, 3 s after start of cue presentation). In the early delay period, we only found the vmPFC. Meanwhile, the late delay period revealed action-specific brain regions, including the l-IPS, l-PMd, and SMA. Thereby, our analysis suggests that the code with which a motor action is initially selected differs from how information is coded further towards motor execution, where the involvement of secondary-motor regions indicate a motor-format coding. This interpretation is corroborated by the results of the cross-regression decoding analysis. In particular, we trained SVRs on the neuronal activation pattern during motor execution and tested for a generalization of these codes in the preceding WM delay. This analysis demonstrates that similar brain-activity patterns in the l-IPS and l-PMd code an anticipated action, as during the actual execution. These results align with a two-stage framework of action planning (Boettcher *et al*., 2021), according to which, after action selection, information about the action outcome is transformed into a motor code during a motor planning stage. The vmPFC likely contributes to action selection (Soon *et al*., 2008), while the l-IPS transforms the intended outcome into a motor code (Gallivan & Culham, 2015; Wong *et al*., 2015; Errante *et al*., 2021; Klautke *et al*., 2023). This motor code could then be stored in the PMd and SMA, where specific motor parameters, such as the to-be used muscle fibres (Mizuguchi *et al*., 2011, 2013, 2014), are selected and maintained in preparation for grip-force execution.

In the following, we will discuss our findings in the temporal sequence from the cue period to the end of the delay period, *i.e.*, until motor execution.

### 4.1 The cue period

During the cue period, no above-chance decoding accuracies were found. Only a small cluster within the vmPFC was revealed when inspecting the data at *p* < 0.001 uncorrected, which however then becomes significant during the early delay period (see discussion in the next session). This finding is consistent with the dynamics of the BOLD response, which is expected to peak at about 3-5 s after event onset, *i.e.*, retro-cue presentation (Martindale *et al*., 2003; Yeşilyurt *et al*., 2008; Hirano *et al*., 2011; Hillman *et al*., 2014). Having found no visual regions suggests that parametric decoding analyses were not influenced by the sensory-features of the visual cue *i.e.*, the grip-force cue, which were experimentally rendered orthogonal to the grip-force intensities. This finding supports the specificity of the main decoding analysis.

The specificity of our SVR decoding analyses is further corroborated by the results of the control analysis, in which decoding of non-maintained grip-force intensities revealed no above-chance decoding at an uncorrected threshold of *p* < 0.001. Finally, label-permutation testing (see *Supplementary Materials*) demonstrates that prediction accuracies of the main analyses are based on the specific parametric coding of grip-force intensities. Together, these null findings and control analyses indicate a high specificity and sensitivity of the reported findings.

### 4.2 The early delay period: action selection in the ventromedial prefrontal cortex

During the early delay period, our main decoding analysis revealed a cluster within the vmPFC; specifically, in Brodmann area 10 (BA-10). This finding is consistent with previous fMRI MVPA studies that implicated the vmPFC in prospective memory tasks (Haynes *et al*., 2007, 2015; Soon *et al*., 2008, 2013; Gilbert, 2011; Momennejad & Haynes, 2012). In these studies, participants were instructed to memorize actions that had to be performed upon the presentation of a relevant cue or event (Haynes *et al*., 2015). Notably, these time-resolved MVPA studies showed that the contribution of the vmPFC to action-selection is limited to an initial processing phase right after cue presentation (Soon *et al*., 2008, 2013).

Alternatively, the vmPFC may have contributed to the maintenance of either a motor intention *i.e.*, the action-selection stage that precedes movement planning (Ruiz *et al*., 2024); also referred to as motor decision stage (Soon *et al*., 2008; Tecilla *et al*., 2022) or a prospective intention *i.e.*, a mental operation regarding the motor details of an intended action (Soon *et al*., 2013). The encoding of prospective intentions is typically tested in prospective memory tasks and operationalized as an abstract, future-oriented memory process involved in the representation and selection of non-motor actions (Soon *et al*., 2013). In contrast, the coding of a motor intention underlies a motor-decision process focused on representing an action outcome and initiating motor planning (Soon *et al*., 2008). While both cognitive processes could explain above-chance decoding in the vmPFC (Soon *et al*., 2008, 2013), the vmPFC’s involvement in coding prospective intentions is unlikely due to a key difference between our delayed grip-force task and traditional prospective memory tasks. Specifically, prospective memory tasks require maintaining intentions while simultaneously performing unrelated secondary tasks during a delay period (Burgess *et al*., 2003; Gilbert, 2011), an absent feature in our task. This distinction complicates direct comparisons with our findings. Conversely, the similarity of our task with those of previous MVPA studies (Soon *et al*., 2008; Ruiz *et al*., 2024) renders above-chance decoding in the vmPFC more likely explained by a contribution to selecting a motor action (Soon *et al*., 2008; Tecilla *et al*., 2022) or the (prospective) motor intentions (Ruiz *et al.,* 2024).

The time-course of the prediction accuracy in the vmPFC, which immediately increases and peaks during the early delay period (similarly to an evoked BOLD response), further suggests that the information processing underlying action-selection begins during the cue period and extends into the early delay period. This finding aligns with our cross-decoding analysis, which indicates stable brain activity patterns in the vmPFC throughout these periods. Consequently, this activity can be viewed as the initial stage of action planning, reflecting the selection of an intended action that is subsequently transformed into a more detailed motor plan (Boettcher *et al*., 2021).

### 4.3 The late delay period: motor planning and grip-force anticipation

Analysis of the late delay period revealed distinct regions compared to the early delay period, namely: the l-PMd, l-IPS and SMA; *i.e.*, regions that are well-known for their role in motor planning.

The most pronounced cluster was within the l-PMd. This finding is in agreement with the results of previous fMRI and TMS studies, which reported the involvement of the PMd in the anticipatory scaling of grip-force (Cole & Rotella, 2002; Chouinard *et al*., 2005; Nowak *et al*., 2009; van Nuenen *et al*., 2012). In an fMRI TMS study, van Nuenen *et al*., (2012) performed univariate analysis, which revealed BOLD signal covariation with the cued grip-force intensities during a delay period. The additional application of low-frequency repetitive TMS (rTMS) over the l-PMd corroborated its involvement in grip-force intensity anticipation (Chouinard *et al*., 2005; Nowak *et al*., 2009; van Nuenen *et al*., 2012). Specifically, rTMS modulated the impact of an incorrect predictive cue on subsequent grip-force execution only when rTMS was applied over the PMd, but not over M1 (Ward *et al*., 2010; van Nuenen *et al*., 2012). This demonstrated that the covariation of the PMd activity with the cued grip-force intensity is essential for accurately anticipating a to-be-executed grip force. A similar contribution of the PMd is suggested by another fMRI study (Mizuguchi *et al*., 2014), which showed brain activity covariation with three imagined grip-force intensities. Our fMRI MVPA study extended these works by employing a multivariate rather than a univariate approach, and by showing parametric modulation of the l-PMd before motor execution. By applying an SVR approach, we directly tested for multivariate codes that map onto a continuous variable, namely, the four anticipated grip-force intensities. Using label-permutation tests, we demonstrated the specificity of the activation patterns within PMd for the particular retained intensity. This code within the PMd is plausible when considering its functional contribution to the representation of anticipating the outcome of a motor movement (Gallivan *et al*., 2011, 2013; Ariani *et al*., 2018; Errante *et al*., 2021; Ariani *et al*., 2022), an effector-independent, action-specific motor plan (Gallivan *et al*., 2011a, 2011b, 2013a, 2011b; Gallivan & Culham, 2015; Ariani *et al*., 2015, 2018), and the maintenance of motor movement plans (Hoshi & Tanji, 2006; van Nuenen *et al*., 2012; Langner *et al*., 2014).

Beneath the l-PMd, activation patterns in the l-IPS were predictive of grip-force intensities during the late delay period. Previous fMRI MVPA studies have consistently shown that the IPS is part of a parieto-frontal network involved in planning grasping actions, even in the absence of fingers pre-shaping (Culham et al., 2003; Frey et al., 2005; Tunik et al., 2005; Davare et al., 2007a; Gallivan et al., 2011a, b; 2013a, b; Ariani et al., 2018; Ruiz et al., 2024; Gallivan et al., 2009; Cavina-Pratesi et al., 2010). These studies have found anticipatory brain-activity in the IPS and decoded movement properties, such as the grip-type during the delay period (Gallivan et al., 2011a, 2013a, 2013b; Ariani et al., 2018; Gallivan et al., 2011b; Ariani et al., 2015). Interestingly, time-resolved decoding analyses have also revealed faster increases of prediction accuracy in the IPS, as compared to effector-specific brain regions (*e.g.*, the SMA) (Gallivan *et al*., 2013; Ariani *et al*., 2018). These findings implicate the IPS into the earlier stages of motor planning, which might even precede motor-preparation (Churchland *et al*., 2010; Shenoy *et al*., 2013; Gallivan & Culham, 2015; Wong *et al*., 2015; Ariani *et al*., 2018; Vyas *et al*., 2020; Ariani *et al*., 2022; Ruiz *et al*., 2024). In particular, the posterior IPS (pIPS) has been shown to represent motor plans in an effector-independent way and to transform intended motor-action outcomes into specific movement plans (Gallivan et al., 2011a, 2011b; 2013a, 2013b; Barany *et al*., 2014; Gallivan & Culham, 2015). This transformation is likely guided by top-down attentional mechanisms, critical for activating and maintaining motor codes in the PMd and selecting to-be executed movements (Schmidt & Blankenburg, 2019; Calton et al., 2002; Beurze et al., 2009; Chang & Snyder, 2010; Szczepanski et al., 2010; Chapman et al., 2011; Gallivan et al., 2011). According to this framework, the pIPS acts as a hub for information processing, where it may transform abstract action outcomes *(i.e.*, the intended action selected in the vmPFC) into detailed movement plans (maintained in the PMd) (Gallivan & Culham, 2015). These findings align with our results, suggesting that the l-IPS may specify intended grip-force intensity in motor plans and support the anticipation and maintenance of motor parameters in the l-PMd by abstractly coding and transforming the intended grip-force intensity into a movement plan (Gallivan et al., 2011a, 2011b; 2013a, 2013b; Gallivan & Culham, 2015).

The functional contribution of the IPS (and PMd) in the maintenance of grip-force intensities is further supported by neurophysiological studies focused on motor-WM (Passingham 1988, 1989; Hoshi & Tanji, 2000; Hoshi & Tanji, 2006; Passingham *et al*., 2007; Nakayama *et al*., 2008; Langner *et al*., 2014; Formica *et al*., 2023). In particular, it has been shown that anatomical and functional connections between the IPS and the PMd facilitate the maintenance of motor information during a delay period (Matelli *et al*., 1986; Johnson *et al*., 1996; Wise *et al*., 1996, 1997; Toni *et al*., 2001; Hoshi & Tanji, 2004, 2006; Pardo-Vazquez *et al*., 2011). While the IPS contributes to the early specification of the to-be executed motor plan (Heed *et al*., 2016; Klautke *et al*., 2023; Ruiz *et al*., 2024), the PMd contributes to the maintenance of effector-specific movement parameters (Passingham 1988, 1989; Kurata, 1993, 1994; Curtis *et al*., 2004; Riehle, 2005; Riehle & Requin, 1989, 1995; Hoshi & Tanji, 2006).

The network of brain regions from which grip-force intensity can be decoded during the delay period showed a third peak in the SMA. The involvement of the SMA in motor preparation has been extensively demonstrated (Marsden *et al*., 1996; Passingham, 1996; Picard & Strick, 1996; Rizzolatti *et al*., 1996; Petit *et al*., 1998; Lee *et al*., 1999; Sreenivasan & D’Esposito, 2019; Bonicalzi & Haggard, 2019), including the preparation of grip-force intensity (van Nuenen *et al*., 2012). It appears plausible that the l-PMd, in interaction with SMA, is involved in generating ready-to-use movement codes (Mizuguchi *et al*., 2011, 2013; van Ede & Nobre, 2023). This is supported by the presence of catch trials requiring rapid and precise grip-force execution, which presuppose the anticipation and maintenance of the intended grip-force in a ready-to-use format (Boettcher *et al*., 2021).

In conclusion, the involvement of pre- and supplementary motor cortices during the late delay period reinforces the view that information is stored in a format - or by the same neuronal populations - as motor movements themselves (Mizuguchi *et al*., 2011, 2013). This hypothesis is corroborated by our cross-regression decoding analysis, which revealed overlapping clusters in the l-IPS and l-PMd during both the late delay period and motor execution. Since executing a motor movement requires transforming action-specific information into a motor code, and a similar code is found during the late delay period, it is likely that information about motor movements was coded well before execution. Interestingly, while generalization to the cue period only showed above-chance decoding in the vmPFC, the late delay period also showed the l-PMd and l-IPS. This difference is likely due to privileged anatomical connections among prefrontal, premotor, and parietal brain regions (Ridderinkhof *et al*., 2004; Huang *et al*., 2019; Löffler *et al*., 2020; Ruiz *et al*., 2024), with the l-PMd serving as a central hub in motor planning and movement preparation (Ruiz *et al*., 2024). This suggests that, once motor decisions are made in the vmPFC, intended grip-force intensities are transformed in the l-IPS and maintained in motor codes of the l-PMd, which is where movement planning occurs. The late increase in prediction accuracy observed in the SMA (at 9 s from cue presentation), combined with below-chance cross-regression decoding accuracy in this area may suggest the SMA’s contribution to initiating motor movements (Gallivan & Culham, 2015; Wong *et al*., 2015; Ariani *et al*., 2018; Vyas *et al*., 2020).

### 4.4 Motor execution

During the motor execution period, our parametric decoding showed above-chance decoding in several brain regions: bilateral primary motor cortices (M1), the l-IPS, l-PMd, SMA and S1 (at *p* < 0.05 FWE-corrected). These results are consistent with previous fMRI studies, showing covariation of M1, SMA, and l-S1 activity with the executed grip-force intensity (Dettmers *et al*., 1995; Thickbroom *et al*., 1998; Dai *et al*., 2001; Ehrsson *et al*., 2001; Cramer *et al*., 2002). The involvement of the M1 and S1 during motor execution, alongside brain-regions found during the late delay period (*i.e.*, the l-IPS, l-PMd, and SMA), is expected. The M1 is well-known for its role in generating motor output (Dechent *et al*., 2004), while S1 plays a crucial role in processing sensory feedback essential for motor control (Wolpert & Flanagan, 2001). Interestingly, while movement-specific information can be decoded from secondary motor regions during both planning and motor execution, above-chance decoding in the M1 is typically limited to the execution period (Ruiz *et al*., 2024). This finding aligns with the results of our main and cross-regression decoding analyses, which did not reveal significant involvement of M1 before motor execution (at *p* < 0.05 FWE-corrected). However, this is only partially consistent with previous fMRI cross-decoding findings (Ariani *et al*., 2018). Specifically, Ariani *et al*. (2018) observed above-chance decoding in both PMd and M1 by cross-decoding between delayed and non-delayed reach-to-grasp tasks. Their delayed condition involved planning and controlling finger kinematics during hand pre-shaping and grasping. Since many neurons in M1 are selective for hand kinematics, this may explain why M1 was engaged in both planning and execution in their study. In contrast, our isometric grip-force task, which does not require finger kinematics planning, could explain the absence of significant M1 involvement until actual execution. The differences between their findings and ours, particularly the lack of above-chance decoding in M1 during motor planning, may be attributed to the distinct task demands in each study (Monaco *et al*., 2020).

In conclusion, results from our parametric analyses corroborate the hypothesis that grip-force anticipation primarily involves the PMd, and not M1 or SMA (van Nuenen *et al*., 2012), indicating a separation of planning and execution functions (Ruiz *et al*., 2024). This differentiation further supports the notion that M1 is more engaged in generating the actual motor output rather than maintaining anticipatory codes (Ruiz *et al*., 2024).

### 4.5 Task-demand dependent WM codes: Prospective *vs* retrospective Working memory

WM representations adapt to the task demands, which employ different neural codes depending on whether the task requires the storage of sensory information (for future comparison with test stimuli) or the transformation of action-specific information into motor movements (Myers *et al*., 2017; Christophel *et al*., 2017). While sensory-WM tasks are used to test the retention of sensory information (Schmidt *et al*., 2017; Uluç *et al*., 2018; Wu *et al*., 2018; Uluç *et al*., 2020), the present study provides critical insights into how WM content is translated into a motor output. In particular, our parametric analyses reveal that information about grip-force intensities is encoded and maintained in distinct brain activity patterns and representational formats across different time-periods of the task. Notably, the presence of similar motor codes during the late delay and motor execution periods supports our hypothesis that intended grip-force intensities were transformed and maintained in a ready-to-use motor format. The presence of WM codes in secondary motor regions corroborates the prospective and action-oriented nature of the undergone task performance.

The action-oriented, prospective nature of our delayed grip-force task allow an interesting comparison with previous research on parametric WM representations, which typically investigate retrospective WM (see Mackey & Curtis, 2017; Christophel *et al*., 2017; van Ede & Nobre, 2023 for further discussion of retrospective and prospective WM functions). Building on the seminal work of Romo *et al*. (1999) in non-human primates, a series of human WM studies have explored different types of parametric WM content. This work has confirmed the relevance of the right inferior frontal gyrus (r-IFG) for the retention of parametric vibratory (Spitzer *et al*., 2010, 2014; Schmidt *et al*., 2017), auditory (Spitzer *et al*., 2012, 2014b; Uluc *et al*., 2018) as well as visual flicker frequency (Spitzer *et al*., 2012; Wu *et al*., 2018). The functional role of the r-IFG in parametric WM is typically interpreted as a reflection of abstract codes of supramodal quantity (Walsh, 2003; Kadosh & Walsh, 2009). Interestingly, we did not find significant above-chance decoding in the IFG (at *p* < 0.05 FWE), even when using more liberal thresholds (at *p* < 0.001 uncorrected). This is a noteworthy difference when aiming to distinguish between retrospective and prospective WM functions. Indeed, it suggests that parametric content representations in the r-IFG are characteristic of retrospective WM, thereby acting as an information buffer for previously perceived data. This information is then maintained in a format suited for abstract processing rather than being directly translated into a motor code. Vice-versa, the reported involvement of the vmPFC in a time-period that precedes transformation into a motor code (and the above-chance decoding in secondary motor regions) may suggest that the vmPFC contributes to the maintenance of parametric, action-specific information for direct translation into motor plans. After translation, this code may be maintained in the PMd. Therefore, our study complements existing literature by revealing that parametric WM codes can retain action-specific information before, and after translation into a motor-code.

## 5 CONCLUSION

In this study, we used MVPA to identify a network of brain regions that encodes parametric WM for the selection, planning and anticipation of to-be executed grip-force intensities. We showed that, after action selection, information on the intended grip-force is transformed into a motor code during a motor planning phase. Specifically, the vmPFC is likely involved in action selection (Soon *et al*., 2008), while the l-IPS contributed to converting this information into a motor code (Gallivan & Culham, 2015; Wong *et al*., 2015; Errante *et al*., 2021; Klautke *et al*., 2023). This motor code is then maintained in the PMd, for the anticipation of grip-force execution in interaction with the SMA. Consistently, cross-regression decoding shows similar activation patterns in the l-PMd and l-IPS during both the late delay period and motor execution, which indicates that action-specific information is retained in a format akin to the motor movement itself (*i.e.*, a motor format; Mylopoulos & Pacherie, 2018).

Our findings align with the view that different types of WM codes are used depending on the task demands (Christophel *et al*., 2017). While much of WM research is focused on sensory-type information in retrospective WM (Schmidt *et al*., 2017; Uluç *et al*., 2018; Wu *et al*., 2018; Uluç *et al*., 2020), the study at hand is an important step to distinguish and characterize how WM content is translated into prospective motor output.

