## Supplementary materials for "Decoding Parametric Grip-Force Anticipation from fMRI-Data"

To test for the specificity of the main decoding analysis, we performed a label-permutation test. Here, the labels of the conditions entered into the SVR were permuted. As previously applied (see Schmidt *et al.*, 2017; Uluc *et al.*, 2020), higher distance of the labelling from the original order should result in reduced decoding accuracies (getting to zero for the highest distance *i.e.*, completely unordered labelling). For all possible permutations, the distance from the rank order was calculated as the sum of the absolute difference of adjacent ranks (*e.g.*, the linear order of grip-force level 1, 2, 3, 4 has a distance of ranks of sum (|1 – 2| + |2 – 3| + |3 – 4|) = 3 and the permuted labelling 2, 1, 3, 4 corresponds to the sum (|2 – 1| + |1 – 3| + |3 – 4|) = 4, resulting in a difference of 1 from the linear order). Thereby, the permutation analysis congregated permutations into four classes of distances from the linear order. For all permutations of labels, the same SVR whole-brain searchlight analysis as in the main decoding was carried out.

Convincingly, when testing in the completely unordered labelling for above chance prediction accuracy in an identical second level ANOVA design, the same t-contrasts across the early and late delay periods revealed no significant clusters throughout the whole brain with identical $p < 0.05$, FWE correction. Even when inspecting this contrast at $p < 0.001$ uncorrected. For illustrative purposes, we display the time-courses of the label-permutation tests with increasing dissimilarity to the original order for the peak voxels of the main analysis in **Supplementary Figure S1 A**. As expected, the time-course of the completely unordered labelling do not show above chance prediction-accuracies throughout all phases of the experimental trials.

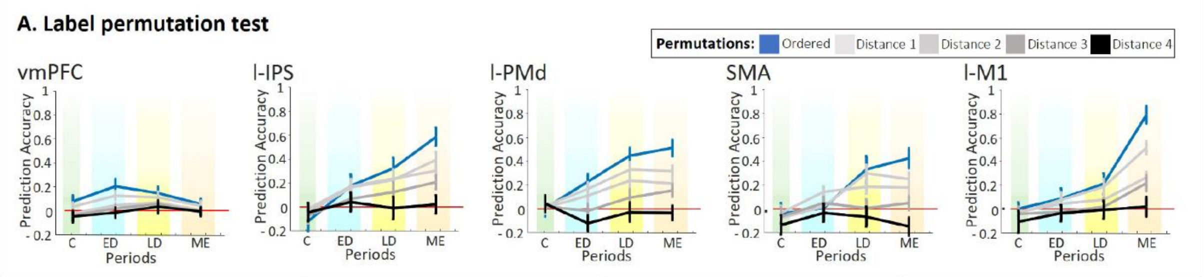


**Supplementary Figure S1 A.** Results of permutation testing in which the same SVR analysis was performed with permuted labels of the data. Prediction accuracy values were extracted from the peak voxels of the five most representative clusters reported in Table 1 (main text). The time-course represents four prediction accuracy values obtained by averaging prediction accuracy values of twelve time-bins in correspondence of the four time-periods (tested in the main analysis). The divergence of the permutations from the linear order of grip-force levels is expressed as distance in rank order. As expected, the divergence from the original order reduces the performance of the SVR

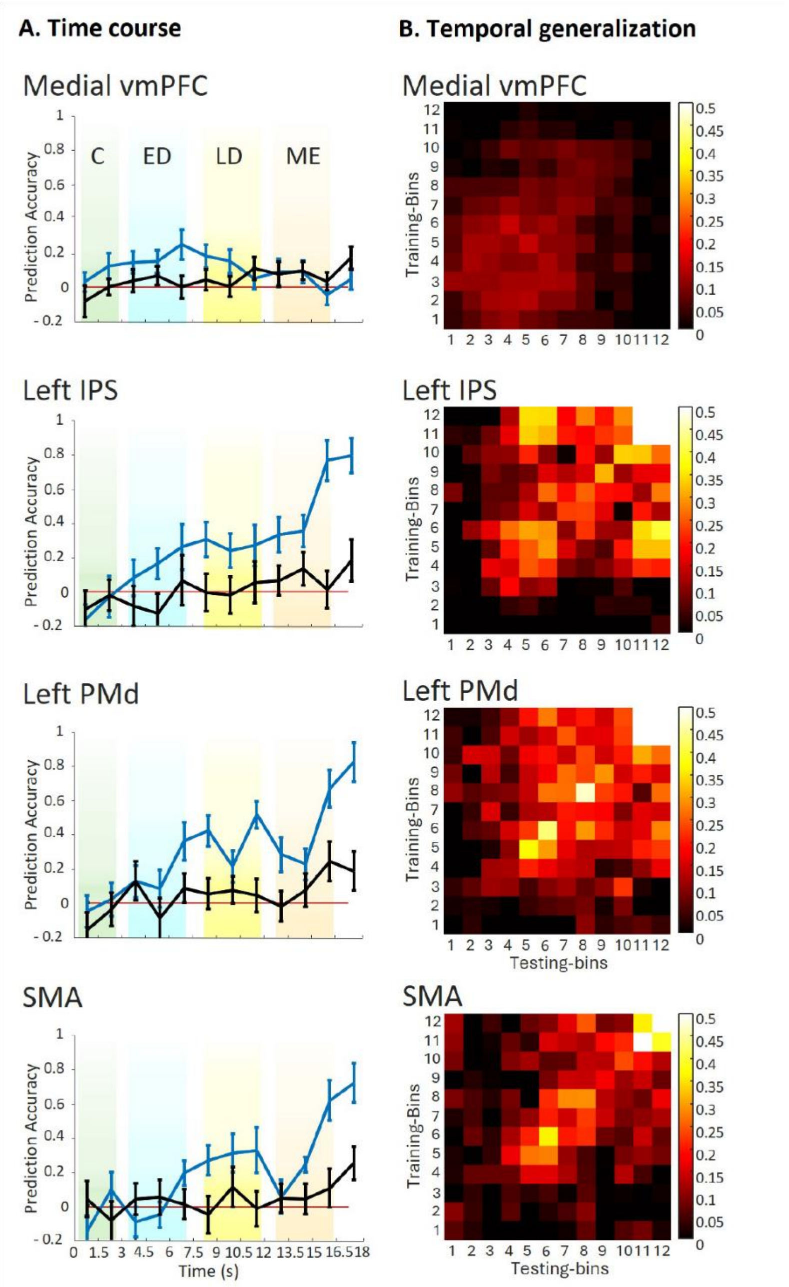


**Supplementary Figure S2 A.** Time-courses of prediction accuracy values (for all 12 time-bins) relative to the main decoding analysis (displayed in blue) and the control analysis (in black), where the non-memorized grip-force level was decoded. Prediction accuracy values were extracted from the peak voxels of the four most representative clusters reported in Figure 3A, and Table 1 (main text). **B.** Temporal generalization matrices, displaying prediction-accuracy values for the four clusters (with lighter colours indicating higher prediction accuracy values). Prediction-accuracy values were extracted from prediction accuracy maps resulting from the whole-brain searchlight cross-regression decoding, where SVRs were trained on all the time-bins (y-axis) and tested on all the time-bins (x-axis), resulting in 144 cross-regression accuracy maps (t1-t12 x t1-t12).

In addition to the performed MVPA, we also explored univariate activation differences during the delay period. We performed two univariate control analyses to test for parametric increases of activity during the delay period, *i.e.*, an analysis based on the same FIR models used for the MVPA, and a second analysis based on classic HRF convolved general linear model (GLM) approach.

At the first level analysis, beta images resulting from the FIR model were normalized and smoothed (with SPMs default of 8 mm FWHM). For each subject, 48 contrast images were computed, *i.e.*, one image for each time-bin and condition; 12 x 4. Contrast images were entered congregated into a flexible factorial second-level design, including a first factor with 4 levels (*i.e.*, one level for force intensity) and a second factor with 12 levels (*i.e.*, one for time-bin). Finally, parametric contrasts were computed based on “Fechner corrected” labels (see main text) for the different periods of the experimental trials. This analysis did not show parametrically modulated activity in the C, ED, LD phases when inspected at $p < 0.05$ FWE corrected.

The classic HRF-convolved analysis confirmed that no relevant parametric modulation of activity was to be found during the delay period. On the first level, the following regressors were modelled as HRF-convolved boxcar functions for the given phases of the trials: the cue period (*i.e.*, t1-t2), delay period (*i.e.*, t3-t8) + parametric modulator, motor execution period (*i.e.*, t9-t11) + parametric modulator, and the motion parameters as regressors of no interest. We used one sample t-test to assess group level effects of first-level contrasts. No significant parametric modulation was found during the delay phase (at $p < 0.05$ FWE-corrected; cluster extent threshold > 10).

Taken together, these univariate control analyses support that the reported MVPA results are not solely based on parametric univariate activation differences during the delay period.
